# Roles for the canonical polarity machinery in the *de novo* establishment of polarity in budding yeast spores

**DOI:** 10.1101/2024.08.29.610423

**Authors:** Benjamin Cooperman, Michael McMurray

## Abstract

The yeast *Saccharomyces cerevisiae* buds at sites pre-determined by cortical landmarks deposited during prior budding. During mating between haploid cells in the lab, external pheromone cues override the cortical landmarks to drive polarization and cell fusion. By contrast, in haploid gametes (called spores) produced by meiosis, a pre-determined polarity site drives initial polarized morphogenesis independent of mating partner location. Spore membranes are made *de novo* so existing cortical landmarks were unknown, as were the mechanisms by which the spore polarity site is made and how it works. We find that the landmark canonically required for distal budding, Bud8, stably marks the spore polarity site along with Bud5, a GEF for the GTPase Rsr1 that canonically links cortical landmarks to the conserved Cdc42 polarity machinery. Cdc42 and other GTPase regulators arrive at the site during its biogenesis, after spore membrane closure but apparently at the site where membrane synthesis began, and then these factors leave, pointing to the presence of discrete phases of maturation. Filamentous actin may be required for initial establishment of the site, but thereafter Bud8 accumulates independent of actin filaments. These results suggest a distinct polarization mechanism that may provide insights into gamete polarization in other organisms.

**SIGNIFICANCE STATEMENT:**

- Dormant budding yeast spores possess a single, stable cortical site that marks the location where polarized growth occurs upon dormancy exit. It was not known how the site forms or which molecules comprise it.
- Using fluorescently tagged proteins in living cells undergoing sporulation, the authors found proteins canonically involved in polarization of non-spore cells arriving at the polarity site in a choreographed manner and required for site function.
- These findings point to a distinct polarity mechanism from non-spore cells and raise new questions about polarity protein interactions with membranes that may be applicable to gametogenesis in other organisms.

## INTRODUCTION

Many of the conserved molecular mechanisms controlling the establishment and maintenance of eukaryotic cellular polarity were first discovered in the budding yeast *Saccharomyces cerevisiae*. For example, the archetypal Rho-like small GTPase controlling polarity, Cdc42, was identified in a genetic screen for temperature-sensitive yeast mutants (Adams *et al*., 1990; Johnson and Pringle, 1990). At high temperatures *cdc42* mutants fail to polarize the cellular actin network to any single point on the plasma membrane (Adams *et al*., 1990) and thus fail to target the trafficking of secretory vesicles (Adamo *et al*., 2001), preventing the selection of a bud site and therefore blocking proliferation.

Subsequent research revealed that in wild-type cells another small GTPase, Rsr1, acts upstream to link recruitment of GTP-bound Cdc42 to a single future bud site on the cell cortex (Park and Bi, 2007; Bi and Park, 2012).

In the absence of Rsr1, yeast cells do break symmetry in a Cdc42-dependent manner, but the site of polarization is random (Irazoqui *et al*., 2003). Budding by wild-type cells is non-random. Haploid and diploid cells both bud at sites adjacent to the prior site of budding, but a diploid cell buds at either of its two poles (Freifelder, 1960). Most *S. cerevisiae* isolates from natural sources are diploid (Peter *et al*., 2018). Yeast meiosis, which is induced in diploid cells by nutrient limitation, typically produces four haploid spores per diploid cell –– two of each mating type –– creating a specialized cell called an ascus (Winge, 1935; Lindegren, 1945). Spores remain mostly dormant until nutrients are provided, whereupon they germinate (Maire *et al*., 2020). To fuse and generate a new diploid, haploid cells must physically contact a haploid partner of the opposite mating type. If germinating spores fail to mate, and bud instead, they are usually able to switch mating types in the cell cycle following their first budding event (Hicks and Herskowitz, 1976). So-called “axial” budding by haploids is thought to promote an efficient return to diploidy by positioning cells of opposite mating type adjacent to each other (Nasmyth, 1983) (Figure 1A). Subsequent fusion between haploid cells creates a diploid zygote. Thus, polarization at non-random cortical locations plays an important role in the budding yeast life cycle.

**Figure 1.**
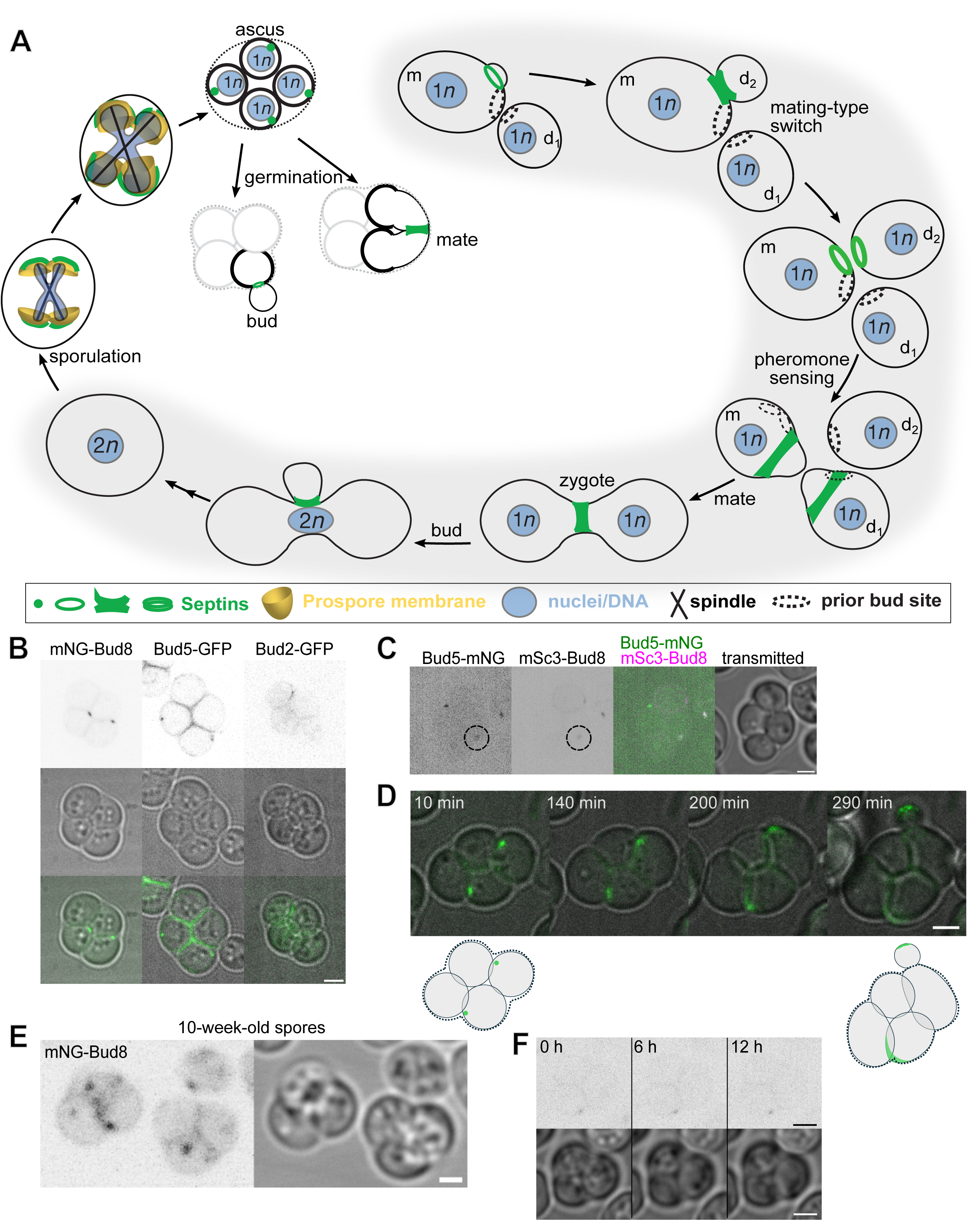
Like the septin Cdc10, Bud8 and Bud5 stably mark the future site of spore polarization upon germination (A) Illustration of septin localization, cell polarization, and morphogenesis during the life cycle of *S. cerevisiae*. Gray background shading highlights events undertaken by vegetative (non-spore) cells in laboratory conditions. Ploidy is indicated by “*n*” values in blue nuclei. “d1” refers to the first daughter cell produced by a mother cell; “d2” is the second; “m” is the mother. The ascus wall is illustrated as a dashed line. The spore wall is shown as a thicker line. Not illustrated here are transient steps in which septins are not discretely localized. Adapted under license CC BY-NC-SA 3.0 from (Spiliotis and McMurray, 2020). (B) Fluorescence or transmitted light micrographs (or merged images thereof) of asci formed by diploid cells with one copy of the gene encoding the indicated polarity protein harboring a fluorescent protein tag. “mNG”, mNeonGreen. Strains were H07248, H07198, and H07227. (C) As in (B) for cells of strain H07284 co-expressing Bud5-mNeonGreen and mScarlet3-Bud8 (“mSc3- Bud8”). The dashed circles surround foci with signal for both tagged proteins. Note that in other asci not shown here, signal strength heterogeneity among foci did not obviously correlate with the presence of the other tagged protein. Scale bar, 2 µm. (D) Wild-type asci (strain H07242) were exposed to rich medium (YPD) for the indicated period of time prior to imaging of mNG-Bud8. Cartoons illustrate cell morphologies and Bud8 localization. Scale bar, 2 µm. (E) Wild-type asci expressing mNG-Bud8 (strain H07248) were imaged following 10 weeks in sporulation medium. Scale bar, 2 µm. (F) A wild-type ascus expressing mNG-Bud8 (strain H07248) in sporulation medium imaged after 0, 6, and 12 hr. Scale bar, 2 µm.

When most spores germinate, they are in immediate proximity to not one but two potential mating partners. When two non-spore (“vegetative” hereafter) cells are placed adjacent to each other in the laboratory, they undergo polarized morphogenesis in response to pheromone signals to extend toward each other and then fuse at the point of contact (Cross *et al*., 1988). By contrast, when a spore mates with one of its meiotic siblings, the two spores do not fuse at the point of contact. We previously discovered that, even before they germinate, *S. cerevisiae* spores are polarized in a manner that, upon germination, promotes their morphogenesis away from the points of contact (Heasley *et al*., 2021).

When sibling spores mate, they do so only after first elongating away from each other, then repolarizing toward each other, creating a zygote with a morphology that is distinct from zygotes formed by non-spore mating (Heasley *et al*., 2021) (Figure 1A). Alternatively, if a spore buds, the bud penetrates the ascus wall (Figure 1A). We speculated that spore polarization prevents buds from becoming trapped within the ascus, and facilitates pheromone sensing prior to mating (Heasley *et al*., 2021). During mating by sibling spores a predetermined polarity program thus initially overrides pheromone signaling. Notably, many natural isolates of *S. cerevisiae* are unable to switch mating types (Fischer *et al*., 2021) and for many isolates competent to switch, mating between sibling spores is the default event upon germination (McClure *et al*., 2018). Hence whereas the mating process in *S. cerevisiae* has been almost exclusively studied using vegetative haploid cells, mating between sibling spores is much more common in nature and likely drove evolution of the underlying polarity pathways, yet it remains largely unexplored.

The mechanism underlying the predetermined spore polarity program was almost entirely unknown. The Cdc10 protein, a member of the highly conserved septin family of proteins that contribute to cell polarization in a wide variety of eukaryotic cell types (Spiliotis and McMurray, 2020), was the only known constituent of the spore polarity site (Joseph-Strauss *et al*., 2007; Heasley *et al*., 2021). During mating and budding by vegetative cells, interplay between septins and Cdc42 is crucial for efficient polarized morphogenesis toward pheromone signal (Kelley *et al*., 2015) and bud site selection (Okada *et al*., 2013), respectively. In each scenario, an upstream signal –– a pheromone gradient or a cortical landmark deposited during prior budding events –– dictates the cortical location where septins assemble (Figure 1A). When they mate, cells fuse at the former sites of pheromone-induced polarization; slow cortical diffusion of residual septins and other polarity proteins biases a diploid zygote when it first buds to polarize near the fusion site (Figure 1A) (Zapanta Rinonos *et al*., 2014).

What recruits a septin to a single site on the spore plasma membrane is not known. The plasma membrane of a spore is formed in a very different way than the plasma membrane of a bud (recently reviewed in (Durant *et al*., 2024; Neiman, 2024)). A double-bilayer membrane, called the prospore membrane (PSM), emerges from a single site at each pole of the meiosis II spindles and elongates in a directional manner around the nucleus, capturing cytoplasm and specific organelles before closing (Figure 1A). This form of cytokinesis is distinct in many ways from cytokinesis in vegetative cells and remains incompletely understood. The outer bilayer of the PSM is degraded after closure, and a stress-resistant cell wall is synthesized around the inner bilayer, which becomes the spore plasma membrane. Septins co-localize with parts of the PSM as it grows (Pablo-Hernando et al., 2008; Fares et al., 1996) and septin mutations result in misdirected PSM growth and spore wall defects (Heasley and McMurray, 2016). Despite differences with budding, PSM biogenesis originating at the spindle poles also involves a form of polarized vesicle trafficking/secretion (Neiman, 1998). Since the extension of the meiosis II spindles during anaphase pushes the spindle poles toward the periphery of the ascus (Figure 1A), we previously proposed that the locations of the spore polarity sites might be explained by polarity factors that persist at the site of PSM origin in the inner bilayer and recruit septins (Heasley *et al*., 2021). Here we explored roles for proteins acting in the canonical yeast polarity pathways in the biogenesis of polarity sites in *S. cerevisiae* spores.

## RESULTS

### The cortical landmark protein Bud8 and the Rsr1 GEF, Bud5, stably mark the spore polarity site

To characterize the molecular composition of the spore polarity spot, we examined cells expressing fluorescently tagged versions of canonical polarity proteins in mature spores. The cortical landmarks that recruit polarity machinery during budding are heavily glycosylated transmembrane proteins that are delivered via post-Golgi secretory vesicles and maintain a history of where polarized secretion occurred. We found the cortical landmark protein Bud8 localized at cortical foci in spores (Figure 1B), similar to the localization of Cdc10 ((Heasley *et al*., 2021) and see below). In vegetative cells, Bud8 links to the cytoplasmic polarity machinery by binding to Bud5 (Kang *et al*., 2004b), the GEF for Rsr1 (Chant *et al*., 1991). We found that Bud5 co-localized with Bud8 to these same foci (Figure 1B and C), while the Rsr1 GAP, Bud2 (Park *et al*., 1993), was not found at these sites in mature spores (Figure 1B). To generate spores for imaging, we used heterozygous diploid cells with only one tagged allele, such that only two of the four spores in an ascus inherit the tagged allele. The fact that Bud5 and Bud8 foci were apparent in two spores per ascus indicates that these signals represent tagged protein that was synthesized after PSM closure.

A germinating spore buds from a single cortical site marked by Cdc10 (Joseph-Strauss *et al*., 2007). When we monitored germination by cells expressing labeled Bud8, we observed that, similarly, buds formed at the location marked by Bud8, indicating that this is the same site (Figure 1D). In every case where we could unambiguously observe it (*n*=28), spores budded from the Bud8-marked site.

Spores can remain nearly dormant for long periods before budding (Maire et al., 2020). Whereas Bud8 is eventually lost from previous bud sites in most vegetative cells (Harkins *et al*., 2001; Kang *et al*., 2022), vegetative cells possess very long-lived transgenerational cortical polarity landmarks with negligible turnover, Rax1 and Rax2 (Chen *et al*., 2000), which interact with Bud8 (Kang *et al*., 2004a). Using established constructs to label the PSM (Nakanishi *et al*., 2007) and Rax1 or Rax2 in vegetative cells (Kang *et al*., 2004a), we could not detect Rax1 or Rax2 at discrete PSM-associated foci (Supplemental Figure 1A and data not shown). However, Bud8 continued to mark foci at the periphery of the ascus for at least 10 weeks following sporulation (Figure 1E), presumably as long as spores remain competent to bud (Maire et al., 2020). Consistent with anchorage in the spore wall, over 12 h cortical Bud8 foci moved a distance of <335 ± 202 nm (n=30) (Figure 1F). Bud8 is unique among the *S. cerevisiae* vegetative landmarks in that polarization always occurs at the site of cortical Bud8 localization, whereas for the other landmarks polarization usually occurs at adjacent locations (Chant and Pringle, 1995). Overall, these observations support a model in which a long-lived canonical polarity landmark, Bud8, is stably deposited at a single site on the cortex of a mature spore and used to mark the site of polarization upon germination.

### Two phases of polarity site assembly

Having identified additional canonical polarity proteins at polarity sites in mature spores, we wanted to better understand how these sites assemble. The exocyst complex is involved in tethering secretory vesicles to sites of docking and fusion during PSM biogenesis (Neiman, 1998; Mathieson *et al*., 2010) as well as polarized exocytosis in vegetative cells (TerBush *et al*., 1996; He and Guo, 2009) and binds directly to multiple canonical polarity proteins, including Cdc42 (Zhang *et al*., 2001). Early exocyst PSM localization (Mathieson et al., 2010) accompanies the initiation of PSM biogenesis at the meiotic outer plaques (MOPs) of the spindle pole bodies (SPBs) (Moens and Rapport, 1971; Peterson et al., 1972; Zickler and Olson, 1975). We previously noted that the deposition of the septin Cdc10 at spore polarity sites could be explained “if septins and/or the exocyst persist on the spore membrane at the former site of contact with the meiotic outer plaque” (Heasley et al., 2021). To test this hypothesis, we monitored localization of septins and canonical polarity machinery, including the exocyst, during PSM biogenesis. PSM closure occurs quickly and is morphologically obvious, so we used it as a point of comparison to determine the relative arrival dynamics of components to the polarity site (Supplemental Movie S1).

Interestingly, following its expected localization to cup-, bar-, and horseshoe-shaped structures associated with the growing PSM (Pablo-Hernando et al., 2008; Fares et al., 1996), the septin Cdc10 appeared at a distinct focus in each spore ∼45 min (median) after PSM closure and then departed ∼120 min thereafter (Figure 2A and see Figure 3). Following PSM closure, the SPBs detach from the spore membranes and move rapidly within the spores, as visualized by co-labeling PSMs and Spc42, a component of the SPB central plaque (Supplemental Figure 1B, Supplemental Movie S1). Polarity sites in mature spores were also distinct from SPBs (Supplemental Figure 1C). Thus, we were unable to directly determine if Cdc10 foci formed at the sites of prior MOP association.

**Figure 2.**
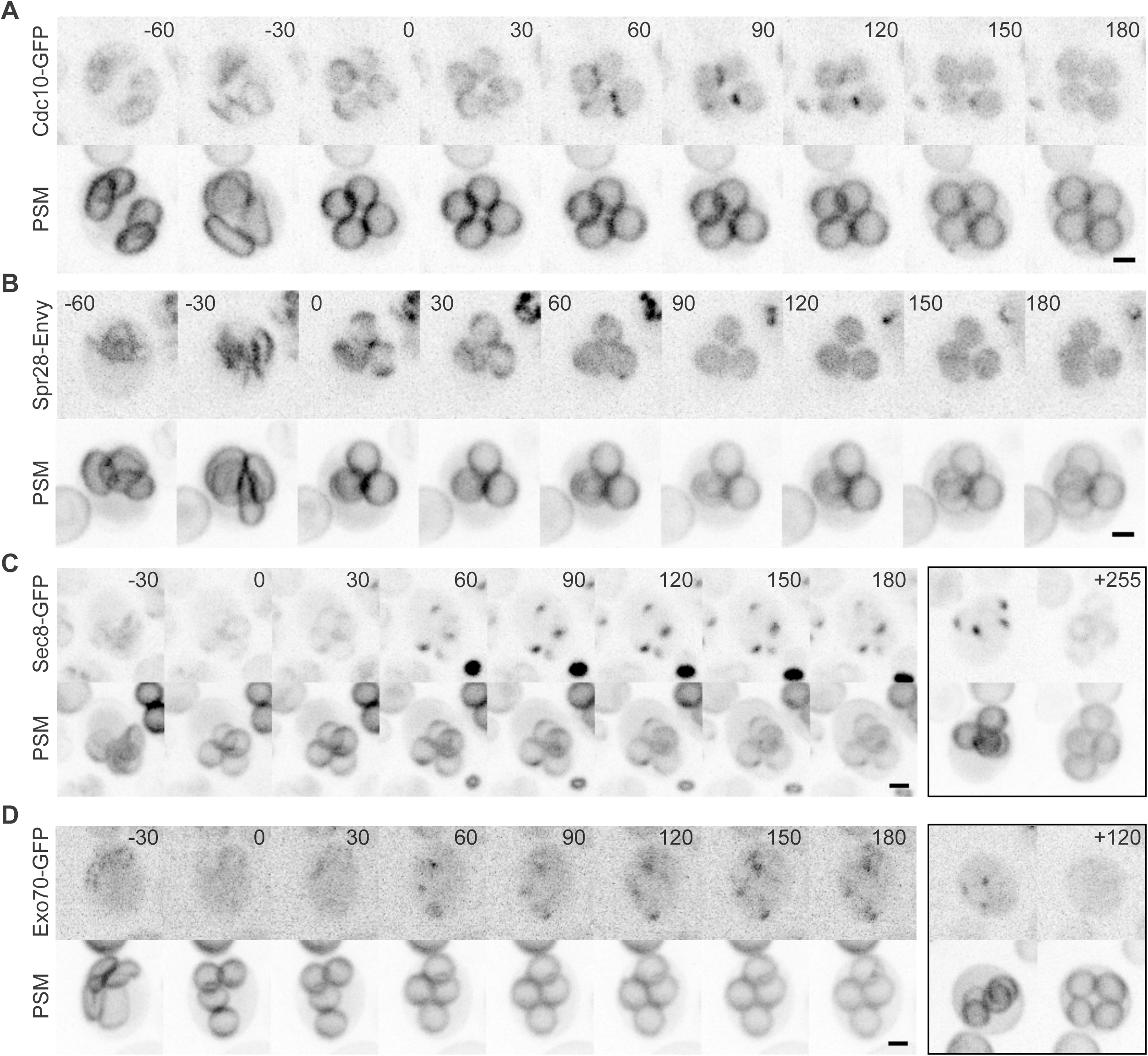
Septin and exocyst localization to a single focus per spore membrane following membrane closure Sporulating cells were imaged at 30-min intervals, with time 0 representing the first time point at which closure of the prospore membrane (“PSM”, marked by an RFP-tagged fragment of Spo20 expressed from plasmid pRS425-R20) had obviously occurred. Scale bars, 2 µm. Cells co-expressed the PSM marker and (A) Cdc10-GFP, strain 132FFBB3; (B) Spr28-Envy, strain 132FFBB3; (C) Sec8-GFP, strain 132FFBB3; or (D) Exo70-GFP, strain H07283. In (C) and (D), the boxed images at right show a different cell from the same experiment in which GFP foci were already visible at closed PSMs at the start of the time course and disappeared during the time course. The numbers in the boxed images indicate how much later, in minutes, the right image was taken relative to the left image.

**Figure 3.**
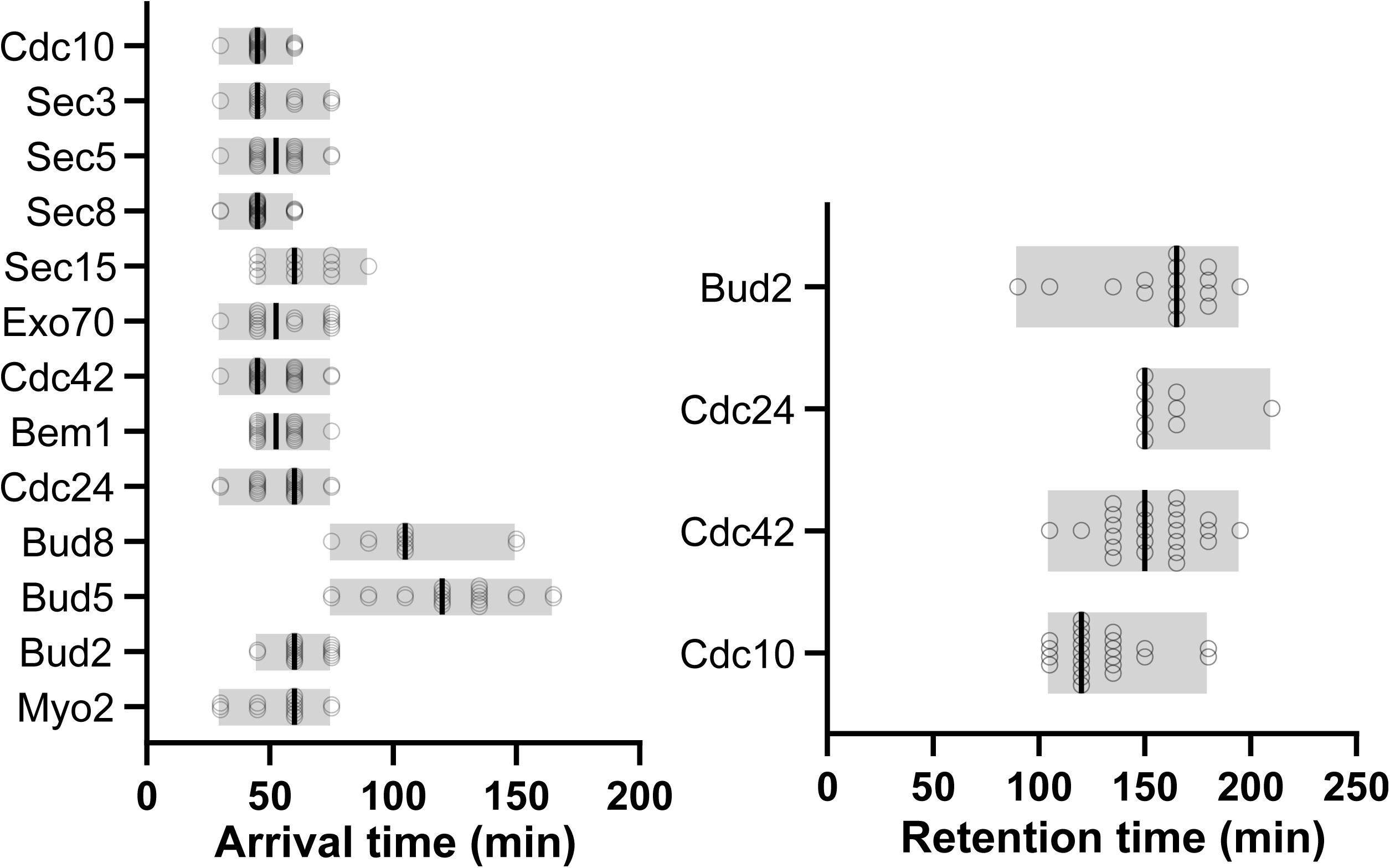
Arrival and retention times for polarity proteins during assembly of the spore polarity site For each of the indicated proteins, “arrival time” was approximated from the time point during experiments like those in Figure 2 when foci of fluorescent signal were apparent at the spore membranes within an ascus (*n ≥* 11 asci per genotype), relative to the time point when PSM closure was first apparent. “Retention time” was calculated similarly (*n ≥* 9 asci per genotype) but from the time point when foci disappeared relative to the time point when foci first appeared. Each value, shown as a circle, represents a single ascus. Lines show medians. Note that Cdc10 returns at some point after departing and, along with Bud5 and Bud8, is found stably associated with the polarity site in mature spores. Data are from strains 132FFBB3, H07191, H06741, H06742, H07182, H07237, H07201, H07202, 01ADAF2E, H07248, H07244, H07227, and H07183 all carrying pRS425-R20 except for strain H07202 which carried pRS426-R20.

Transient Cdc10 foci were visible in all four spores within an ascus formed by diploid cells heterozygous for the *CDC10-GFP* allele, indicating that the same Cdc10 molecules synthesized prior to PSM closure were used to assemble these foci. The septin structures associated with the growing PSM include the sporulation-specific septin Spr28 (Pablo-Hernando et al., 2008; De Virgilio et al., 1996) (Figure 2B). Spr28 was previously found in a yeast two-hybrid screen to interact with a canonical polarity protein (Drees et al., 2001). However, Spr28-Envy was not found at any PSM foci following PSM closure (Figure 2B), suggesting that the Cdc10-containing foci at spore polarity sites are composed of distinct septin complexes from those involved in PSM biogenesis.

As with Cdc10, we saw localization of GFP-tagged molecules of the exocyst subunits Exo70 and Sec8 to a single cortical focus in each spore ∼45 min after PSM closure (Figure 2C,D). These foci persisted throughout the duration of our imaging (∼4 h) but observations of mature spores demonstrate that these foci eventually disappear (Figure 2C,D). We repeated this experiment using labeled versions of other exocyst subunits (Sec3, Sec5, Sec15) and saw similar results (Supplemental Figure 1D, Figure 3).

Given the role of the exocyst in the targeting of polarized trafficking, we reasoned that the master polarity regulator, Cdc42, would also localize to such foci. During PSM biogenesis, GFP-tagged Cdc42 was found rather diffusely on the growing membrane. GFP-Cdc42 signal then accumulated ∼45 min after PSM closure at a single focus visible in each of the four spore membranes within an ascus (Figure 3, 4A). Cdc42 persisted at these foci for ∼150 min, then departed and did not return during the remainder of sporulation (Figure 3). The duration of Cdc42 focus persistence was similar to that of Cdc10 (Figure 3) though due to the limited temporal resolution of our imaging (15-min timepoints) we may have missed subtle differences.

The scaffolding protein Bem1 (Butty *et al*., 2002; Kozubowski *et al*., 2008) is thought to stabilize the assembly of polarity complexes on membranes where it recruits Cdc24, a GEF for Cdc42 (Peterson *et al*., 1994; Zheng *et al*., 1995; Ito *et al*., 2001). We found that Bem1 and Cdc24 both formed foci at sites on spore membranes following similar dynamics, arriving ∼50 min after closure (Figure 3, 4B). Cdc24 remained at these sites for ∼150 minutes (Figure 3) while Bem1 remained for the duration of our imaging, though it is not present in mature spores. Unexpectedly, Bud8 and Bud5 arrived later (>100 min after PSM closure) than the other canonical polarity proteins (Figure 3, 4D and E). While Bud2 was not found at polarity sites in mature spores, it was detected as foci appearing ∼60 min after PSM closure (Figure 3, 4F), which more closely matches the arrival times of Cdc42, Cdc24, and Bem1. Bud2 foci at spore membranes persisted for ∼165 min (Figure 3). These observations support a model in which there are multiple phases of polarity site assembly that are distinguished by the presence of canonical polarity proteins arriving and departing in groups that mostly correspond to the functions of these proteins in vegetative cells, with some unexpected differences. Overall, the polarized vesicle trafficking machinery and the master polarity GTPase (Cdc42) arrive first, recruiting the associated regulatory machinery to generate positive feedback loops necessary for stable polarization, eventually leading to the recruitment of Bud8, Bud5, and Cdc10, which remain at the polarity site until germination.

### Mutating canonical polarity genes disables the spore polarity site

We next examined the requirement for canonical polarity factors in assembling a functional spore polarity site, using mScarlet3-Bud8 and Bem1-GFP as markers. Cells lacking *BUD2*, *BUD5*, or *RSR1* formed Bud8-marked polarity sites but, unlike wild-type cells, upon germination mutant cells often polarized and budded at other cortical locations (Figure 5, Supplemental Movies S2-S5). It remains unclear whether this phenotype is due to defects arising during the assembly of the site, its usage, or both. These findings demonstrate roles for the canonical polarity machinery in establishing and/or functionalizing a specific cortical site in the spore membrane that is built shortly following PSM closure and is used upon germination to drive cellular polarization.

**Figure 4.**
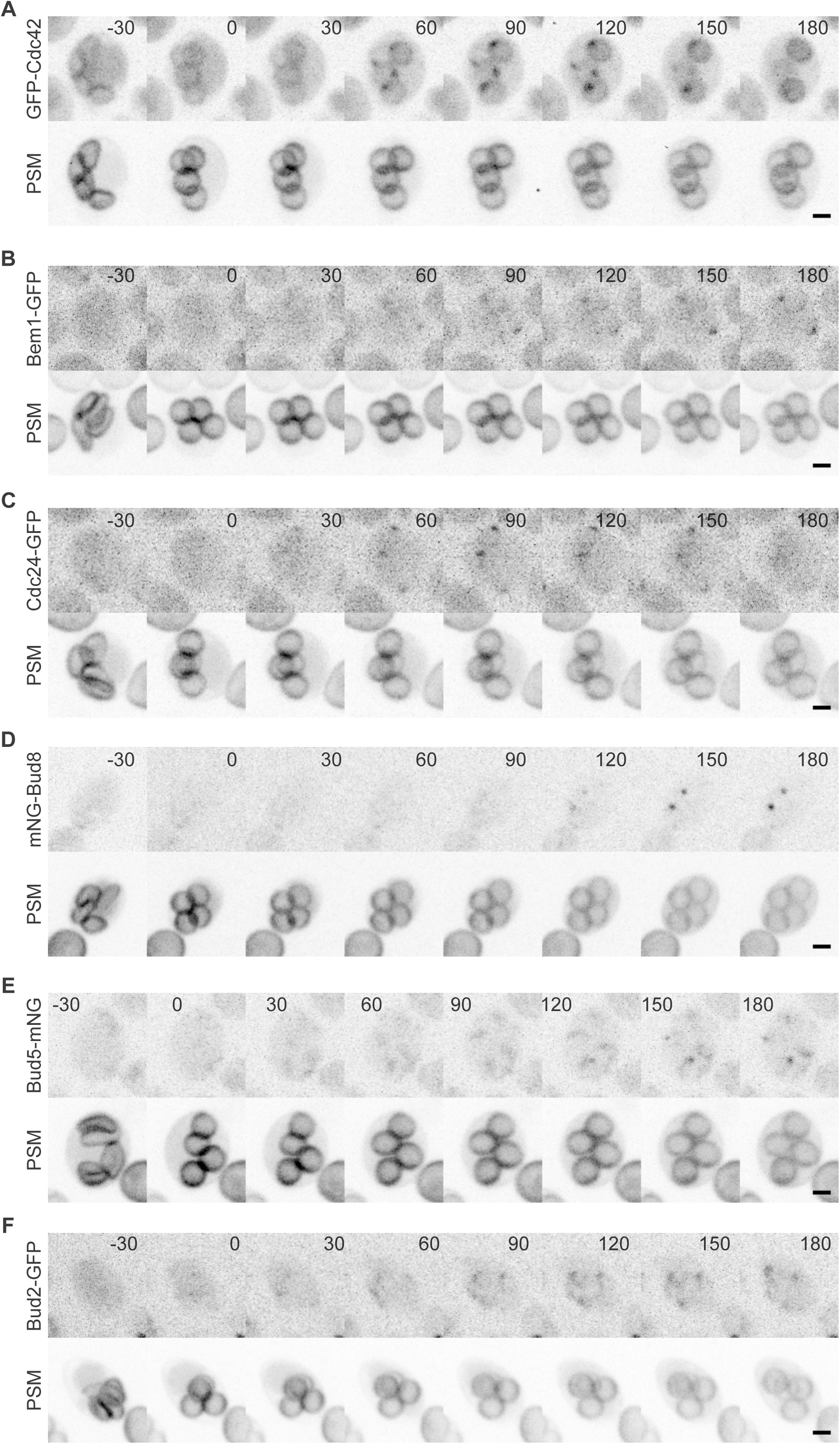
Localization of canonical cell polarity proteins to a single focus per spore membrane following membrane closure As in Figure 2, sporulating cells were imaged at 30-min intervals, with time 0 representing PSM closure. Scale bars, 2 µm. All strains expressed the same PSM reporter. (A) Strain H07201 with plasmid pRS425-R20, expressing GFP-Cdc42. (B) Strain H07202 with plasmid pRS426-R20, expressing Bem1- GFP. (C) Strain 01ADAF2E with plasmid pRS425-R20, expressing Cdc24-GFP. (D) Strain H07248 with plasmid pRS425-R20, expressing mNeonGreen-Bud8. (E) Strain H07244 with plasmid pRS425-R20, expressing Bud5-mNeonGreen. (F) Strain H07227 with plasmid pRS425-R20, expressing Bud2-GFP. Scale bars, 2 µm.

**Figure 5.**
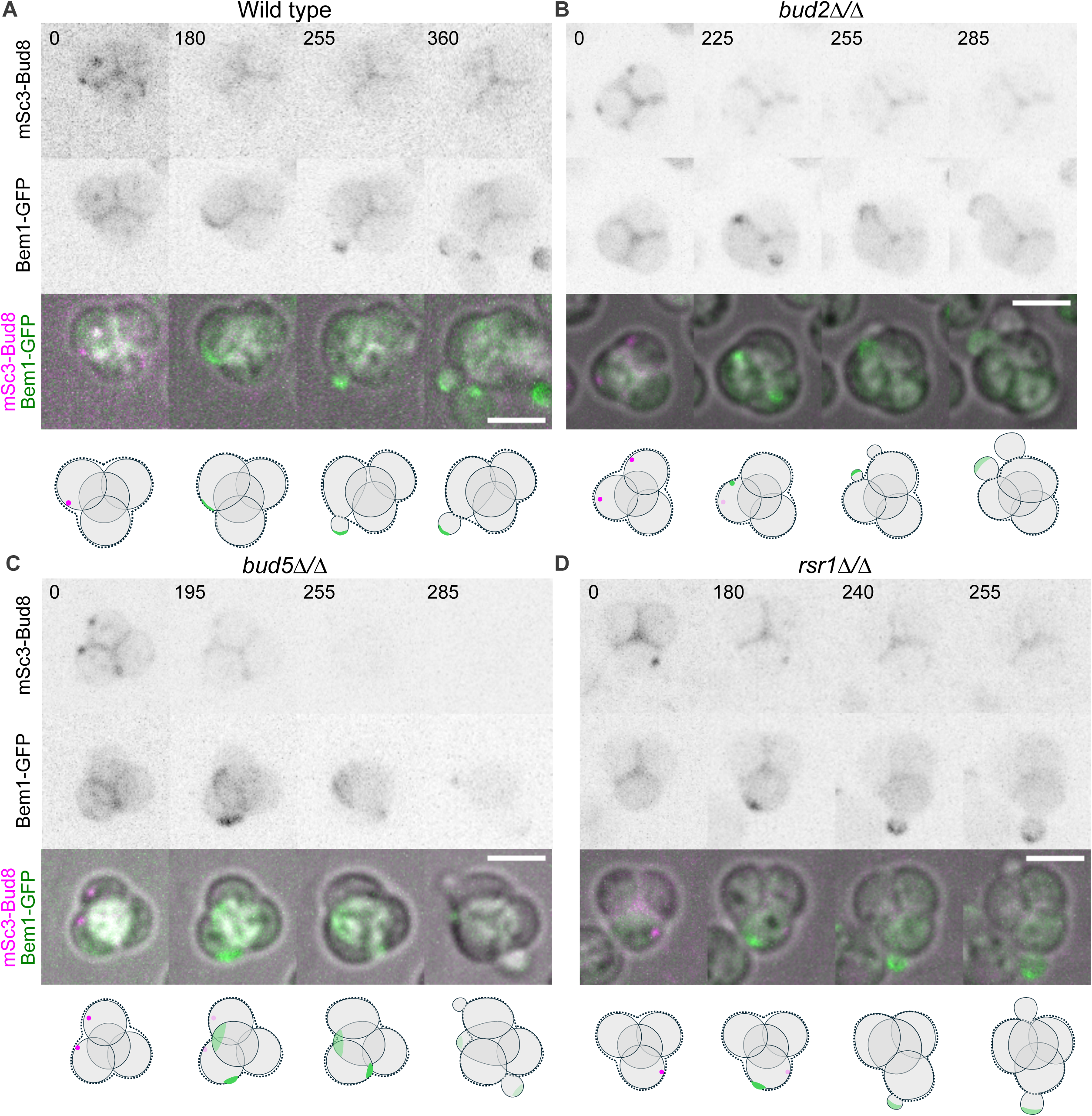
Mutating canonical polarity genes unlinks polarization during germination from the Bud8-marked site As in Figure 1D, asci were exposed to rich medium (YPD) to induce germination for the indicated period of time prior to imaging. Cells of the indicated genotypes co-expressed mScarlet3-Bud8 and Bem1-GFP. Cartoons illustrate cell morphologies and the relevant localization of Bud8 and Bem1. Scale bars, 5 µm. (A) Wild-type ascus of strain H07280. (B) *bud2Δ/Δ* ascus of strain H07282. (C) *bud5Δ/Δ* ascus of strain H07281. (D) *rsr1Δ/Δ* ascus of strain H07283.

### Polarized trafficking in the assembly of the polarity site

Both active vesicle trafficking and passive, diffusion-mediated events are known to facilitate the localization of polarity factors to specific cortical locations on the yeast plasma membrane (Brauns *et al*., 2023). The appearance of exocyst at foci on spore membranes following PSM closure is consistent with polarized vesicle trafficking, typically associated with transport via myosin motors like yeast Myo2 (Johnston *et al*., 1991). In vegetative cells, Myo2 has been shown to associate with the exocyst, facilitating the delivery of secretory vesicles to targeted membranes (Jin et al., 2011). We observed Myo2 arriving at cortical foci ∼60 min after PSM closure (Figure 3, 6A). Similar to the exocyst components, Myo2 signal remained present for the duration of our imaging but was not present in mature spores. In cells expressing both Myo2-GFP and mScarlet3-Bud8, we observed that Myo2 foci formed first, followed by the later appearance of Bud8 foci at the same site (Figure 6B).

**Figure 6.**
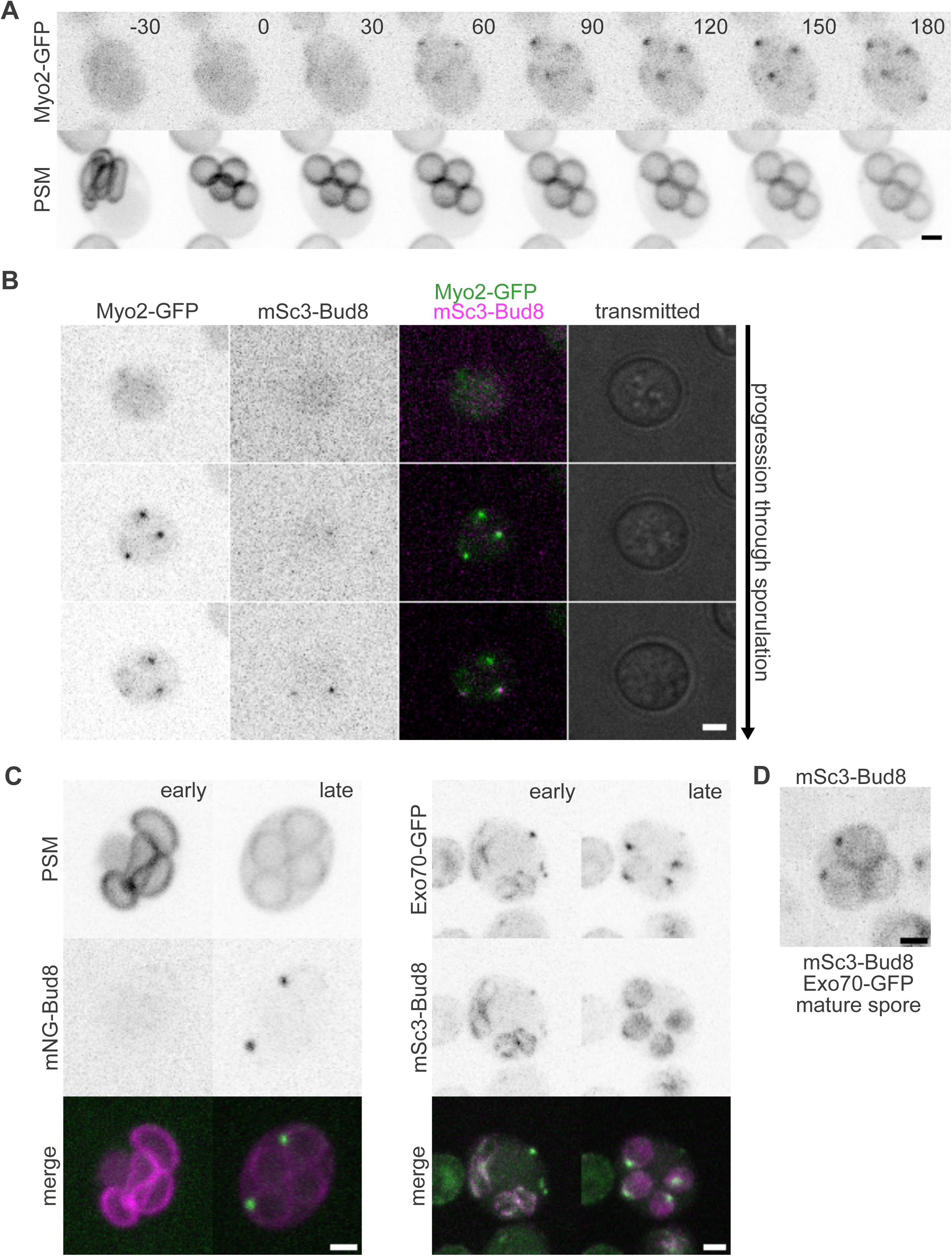
Localization of the motor protein Myo2 to the spore polarity site and Bud8 localization defects upon tagging exocyst (A) As in Figure 2, sporulating cells were imaged at 30-min intervals, with time 0 representing PSM closure. Shown is a representative ascus of strain H07183 carrying plasmid pRS425-R20. Scale bar, 2 µm. (B) Asci of strain 7FAEB7DC, in which Myo2 is tagged with GFP and Bud8 is tagged with mScarlet3 (“mSc3”), were imaged at various timepoints during sporulation. The color image shows an overlay of Bud8 and Myo2 signals. Scale bar, 2 µm. (C) A representative ascus of strain H07248 carrying plasmid pRS425-R20 (left) or C203ACF0 (right) imaged at a time point just before (“early”) and after PSM closure (“late”), showing Bud8 tagged with mNeonGreen (“mNG”) or mScarlet3 and the PSM reporter or GFP-tagged exocyst subunit Exo70. Scale bars, 2 µm. (D) A representative ascus of strain C203ACF0 containing mature spores, showing clear Bud8 foci at spore membranes despite earlier mislocalization throughout spore membranes shortly after PSM closure, as shown in (C). Scale bar, 2 µm.

To determine if Bud8 co-localizes with exocyst foci, we co-labeled Exo70-GFP in cells also expressing mScarlet3-Bud8. To our surprise, the combination of these tags caused a striking mislocalization of mScarlet3-Bud8 throughout the growing PSMs, rather than only to a single focus following PSM closure (Figure 6C, Supplemental Movie S6). Localization of mScarlet3-Bud8 to all four PSMs within a developing tetrad in tagged-exocyst cells carrying only one copy of the *mScarlet3-BUD8* gene also contrasts with the appearance of tagged Bud8 foci in only two of four spores in the absence of tagged Exo70 (Figure 6C, Supplemental Movie S7). In the latter cells, the two spores with tagged Bud8 foci presumably represent the two spores that inherited the tagged *BUD8* allele, suggesting that normally only Bud8 synthesized following PSM closure is trafficked to the polarity site. Furthermore, we noticed that while Bud8 foci did eventually appear in cells with tagged exocyst, the timing was delayed relative to cells without labeled exocyst, and the foci appeared in only two of the four spores in an ascus (Figure 6D). Since exocyst is required for the earliest steps in PSM biogenesis (Neiman, 1998), it would have been challenging to deliberately perturb exocyst function and look for effects on Bud8 localization without completely disrupting spore membranes. Instead, we were able to exploit unexpected, subtle defects in exocyst function accompanying fluorescent tagging to demonstrate a requirement for exocyst in the proper dynamics of Bud8 delivery to the spore polarity site.

### Roles for actin in the polarized trafficking of polarity site components

The apparent involvement of Myo2 in the assembly of the polarity site (Figure 6A) is consistent with actin-mediated transport. It has been shown that filamentous actin structures play a role in the delivery of precursor elements during PSM biogenesis (Taxis *et al*., 2006), though specific details remain unknown. Similar to previous reports (Morishita and Engebrecht, 2005; Taxis *et al*., 2006), when we used Lifeact-GFP to label actin structures in sporulating cells, we noticed that although cables were present in meiotic cells, only actin patches could be detected after PSM closure (Supplemental Figure 2A, Supplemental Movie S8). A new, more sensitive Lifeact-3xmNG fusion (Wirshing and Goode, 2024) did not provide clearer results (Supplemental Figure 2A). Bright patches might limit our ability to see dimmer cables, so we looked instead for indirect evidence of actin cables. An actin-nucleating formin is among the MOP proteins in sporulating cells of the filamentous fungus *Ashbya gossypii*, a close relative of *S. cerevisiae* (Kemper et al., 2011). However, we failed to detect discrete fluorescent signal in sporulating *S. cerevisiae* cells expressing tagged versions of the budding yeast formins Bnr1 and Bni1 (Supplemental Figure 2B).

We tried to test formin function in spore polarity site assembly but diploid cells lacking one or both copies of *BNI1* failed to sporulate: we could not find a single ascus in sporulation cultures. The Bud6 protein interacts with Bni1 and contributes to actin nucleation (Graziano *et al*., 2011). Bud6-GFP formed bright foci following PSM closure but these often appeared to be outside of the PSMs and did not localize in any consistent pattern in mature spores (Supplemental Figure 2C). Similarly, Arp2, a marker of actin patches (Winter *et al*., 1997), and Ede1 and Abp1, markers of distinct phases of endocytosis (Layton *et al*., 2011), were scattered around the spore membranes following PSM closure but amidst the multitude of dynamic signals we were unable to detect specific foci that might represent polarity sites (Supplemental Figure 2D). These signals presumably represent other, more general actin-dependent events that are likely involved in spore wall assembly (Morishita and Engebrecht, 2005; Taxis *et al*., 2006) and/or disassembly of the plasma membrane of the diploid mother cell.

We took two approaches to perturb actin-based trafficking and monitor assembly of the spore polarity site. In vegetative cells carrying the temperature-sensitive *cdc42-1* allele, the actin cytoskeleton is depolarized within 1 hr of incubation at 33°C (Adamo *et al*., 2001). The original *cdc42-1* allele carries multiple mutations but only a single coding change relative to the genome reference strain (Miller and Johnson, 1997). We found that the G142S substitution alone was sufficient to confer a temperature-sensitive phenotype (Figure 7A). However, Exo70-GFP localized normally to the spore polarity site in *cdc42(G142S)* cells shifted to 37°C on the microscope stage just after PSM closure (Figure 7B). We verified the efficacy of the temperature shift using vegetative cells carrying the *cdc12-6* allele

**Figure 7.**
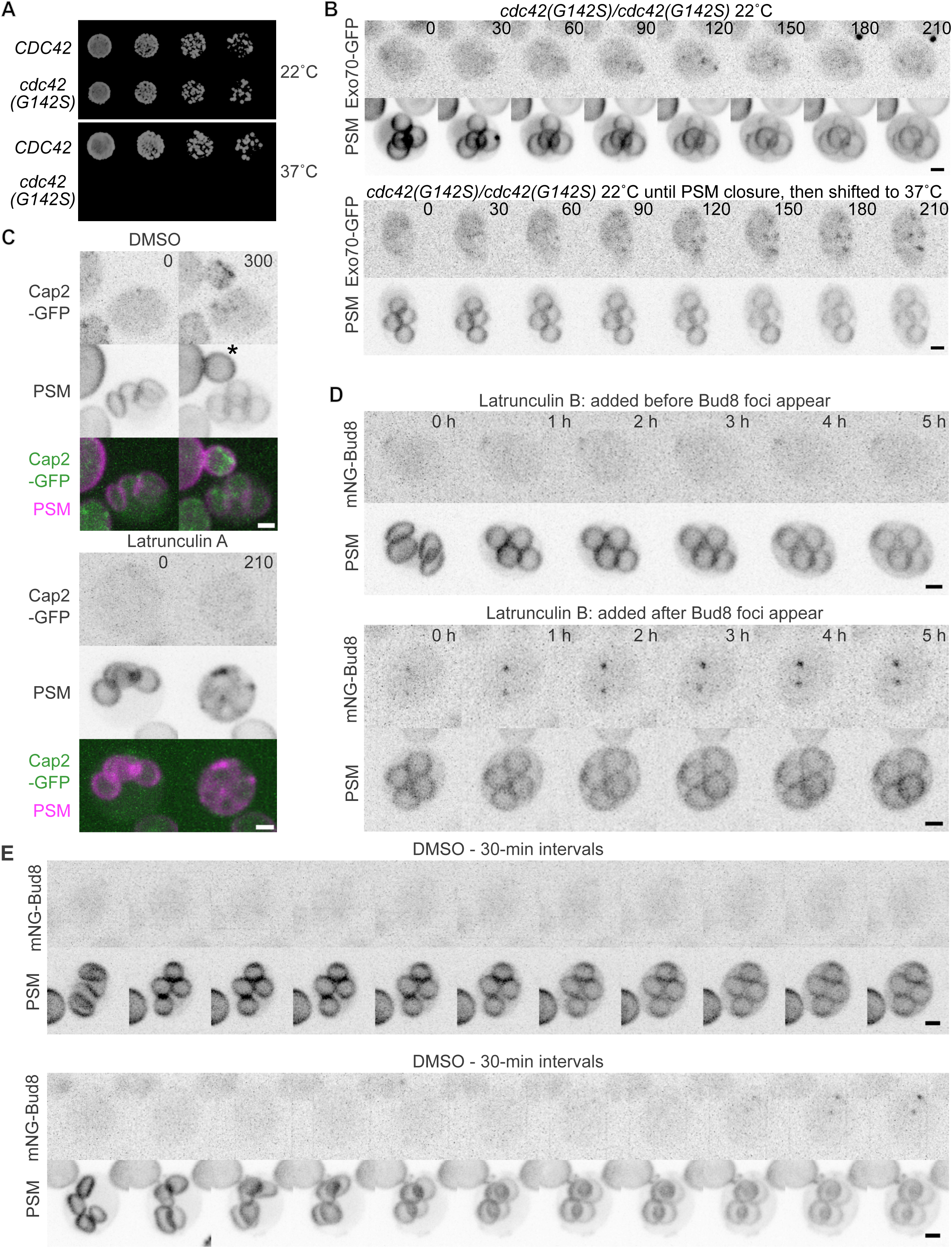
Following initial recruitment, polarity protein accumulation at the spore polarity site does not require actin filaments (A) Five-fold serial dilutions of cells of strains H07237 (“*CDC42*”) and 0C39D4FC (“*cdc42(G142S)*”) were spotted on rich agar medium and incubated at the indicated temperature for four (top) or three (bottom) days prior to imaging. (B) Sporulating cells of strain 0C39D4FC carrying plasmid pRS425-R20 that co-express the PSM reporter and a GFP-tagged version of the exocyst subunit Exo70 were imaged at 30-min intervals for 210 minutes. The top series shows a representative ascus maintained at 22°C throughout. The bottom series shows a representative ascus that was shifted to 37°C just after PSM closure. Scale bars, 2 µm. (C) Sporulating cells of strain A76EDCB8 carrying plasmid pRS425-R20 that co-express the PSM reporter and a GFP-tagged version of the actin patch marker Cap2 were exposed to 1% DMSO (as a solvent control) or 200 µM Latrunculin A dissolved in DMSO. Numbers indicate the time (in minutes) between addition of DMSO or Latrunculin A and imaging, not counting ∼10 min required to mount the cells for imaging. A representative ascus is shown for each condition. Scale bars, 2 µm. The asterisk indicates a bud produced by an adjacent vegetative cell. (D) Sporulating cells of strain H07248 carrying plasmid pRS425-R20 that co-express the PSM reporter and mNeonGreen-tagged Bud8 (“mNG-Bud8”) were exposed to 200 µM Latrunculin B and imaged at 1- hr intervals. As indicated, the two asci shown are representative of what was observed when Bud8 foci had not yet appeared at the time of Latrunculin addition versus when Bud8 foci were already visible. A Cap2-GFP-expressing ascus treated in the same way is shown in Supplemental Figure 2E. Scale bars, 2 µm. (E) As in (D), cells of strain H07248 carrying plasmid pRS425-R20 undergoing sporulation were exposed to 1% DMSO. Two asci are shown, with the top representing what was most commonly seen, and the bottom showing a rare occasion when Bud8 foci formed. Scale bars, 2 µm.

(Dobbelaere *et al*., 2003) with a GFP tag fused to the mutant Cdc12: septin rings disappeared within ∼30 minutes of 37°C shift (Supplemental Figure 2F).

We found similar results when we pharmacologically ablated filamentous actin in sporulating cells using latrunculin, an actin-binding toxin produced by sea sponges (Spector *et al*., 1983; Coué *et al*., 1987).

We used GFP-tagged Cap2, a subunit of the heterodimeric F-actin capping protein complex, to label actin patches as a reporter of F-actin (Waddle *et al*., 1996). Addition of latrunculin A (Lat A) at 200 µM blocked full extension and closure of PSMs that had not yet closed prior to Lat A addition, and PSMs that had already closed began to break down within the same time frame as Bud8 foci appear in wild-type cells (Figure 7C), precluding assessment of polarity site assembly. It is not clear why in other studies the same Lat A concentration did not have such severe effects (Taxis *et al*., 2006) but given the evidence of robust endocytic activity following PSM closure these effects may reflect requirements for endocytosis in PSM persistence. Latrunculin B (Lat B) is less potent at disassembling yeast F-actin (Irazoqui *et al*., 2005). At 200 µM, addition of Lat B led to the full ablation of Cap2-GFP foci, but PSMs did not break down (Figure 7D, Supplemental Movies S9-10). In spores where Bud8 foci had just begun to appear, signal at these foci continued to increase following Lat B addition (Figure 7D, Supplemental Movie S10). While we were occasionally able to see polarity sites assemble in DMSO-treated cells (Figure 7E, Supplemental Movie S9), DMSO strongly reduced overall sporulation efficiency and is known in other contexts to alter membrane properties (Notman *et al*., 2006; Gurtovenko and Anwar, 2007; Gironi *et al*., 2020). Consequently, we cannot exclude the possibility that DMSO itself also inhibits initial polarity site assembly. Thus, while our data suggest that filamentous actin may be essential to the initial process of polarity site assembly, once a site is assembled, subsequent trafficking of Bud8 to the site does not directly require filamentous actin.

## DISCUSSION

### Non-canonical roles for canonical polarity proteins

The new results we obtained in this study support the model we previously proposed (Heasley *et al*., 2021), with a few surprises (Figure 8). One surprise was that the cortical landmark Bud8, which is not especially long-lived in vegetative cells, persists for weeks at the spore polarity sites. By contrast, Rax1 or Rax2, the long-lived vegetative landmarks, were not found at these sites. Bud5 persistence with Bud8 matches what is seen in vegetative cells, where Bud8 and Bud5 can be found at the most recent bud site but not at older sites (Harkins *et al*., 2001; Marston *et al*., 2001). Bud2 disappears from the bud neck prior to the completion of cytokinesis (Park *et al*., 1999), similar to the absence of Bud2 at the polarity site in mature spores.

**Figure 8.**
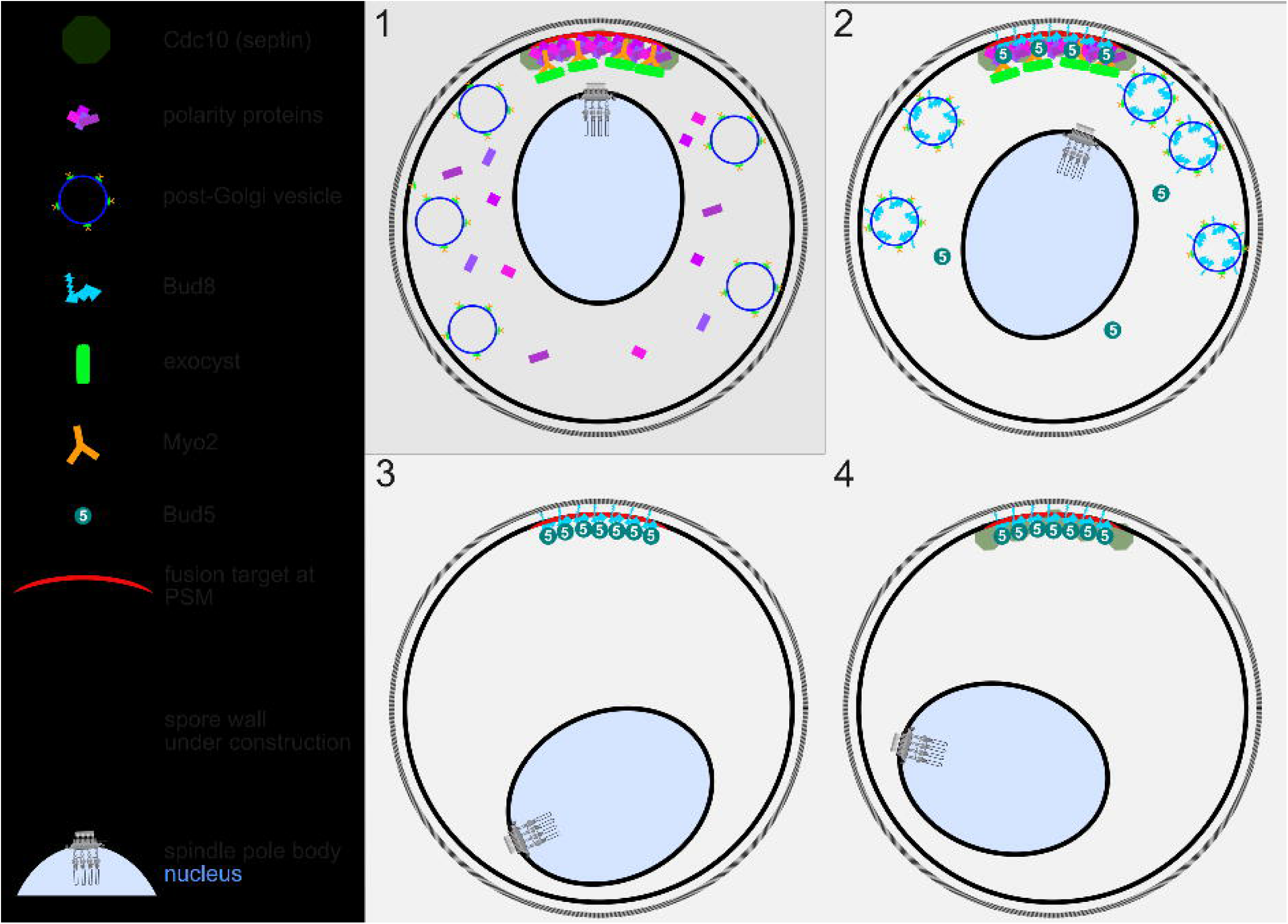
Model of spore polarity site assembly in two phases Illustrated are two phases of spore polarity site assembly, both occurring following PSM closure. During the “set up” phase (1) post-Golgi vesicles bound to Myo2 fuse at the PSM origin, which is marked by an unknown fusion target, potentially a specific phospholipid. Canonical polarity proteins (including Cdc24, Cdc42, Bem1, and Bud2) begin to accumulate at the polarity site as feedback loops develop. Three steps (2-4) comprise the “completion” phase; while they are shown as discrete steps, they likely overlap. (2) Feedback loops enforce polarity at the site, directing the fusion of newly generated vesicles which now contain Bud8. Bud8 is embedded in the membrane at the polarity site where its position is stabilized through contacts with the newly synthesized spore wall. Soluble Bud5 begins to accumulate at the polarity site through interactions with Bud8. Bud5 triggers a gradual breakdown of the polarity feedback loops, perhaps by triggering Bud2 exit (not shown). A “dilution” effect as new membrane is delivered may contribute to weakening of feedback loops; we cannot rule out contributions of other negative regulators. In step 3, the canonical polarity proteins have left the site. (4) The return of Cdc10 marks the final step of the completion phase, resulting in a spore with a mature polarity site.

Two distinct mechanisms of Cdc42 activation have been identified in vegetative cells, though the details remain somewhat controversial. Some studies suggest that in early G1, pre-START, Bud3 acts as the Cdc42 GEF, whereas just after START, Cdc24 is responsible for Cdc42 nucleotide exchange, scaffolded by Bem1 (Kang *et al*., 2014; Miller *et al*., 2019). On the other hand, there is evidence of pre-START activation of Cdc42 via Bem1 and Cdc24 in daughter cells, with the Cdc42 effector Ste20 acting unlike other effectors (including the exocyst subunit Exo70) in co-localizing with Cdc42 in pre-START G1 (Moran *et al*., 2019). As we did not detect any discrete Bud3 localization during sporulation (Supplemental Figure 3A), we interpret the arrival of Cdc24 and Bem1 at about the same time as Cdc42 as evidence that Cdc24 acts as the relevant GEF for the spore polarity site. We did not find any evidence for discrete localization of Ste20 during or upon completion of sporulation (Supplemental Figure 3B); conversely, we did find Exo70 arriving at the site of spore polarity site assembly at about the same time as Cdc42 and its regulators (Figure 2D). Thus, our findings during sporulation may represent an additional, previously unknown mechanism of Cdc42 regulation.

We found evidence of multiple phases of biogenesis of the spore polarity site following PSM closure (Figure 8). In the first, exocyst, the septin Cdc10, and the canonical polarity proteins arrive. Bem1 can directly bind exocyst (France *et al*., 2006) and so can Cdc42 (Zhang *et al*., 2001). A recent report also provided evidence that fission yeast septins, including the Cdc10 homolog Spn2, are able to directly interact with exocyst subunits (Singh and Kruger, 2009). Thus by virtue of direct interactions with Myo2 (Jin *et al*., 2011), exocyst could deliver any of these polarity proteins to the spore polarity spot without needing to also transport vesicles, but we assume that vesicles are likely associated with exocyst and Myo2 at this point (Figure 8). Bud8 and Bud5 then arrive, and shortly after, Cdc10 and the other polarity components leave. Exocyst and Myo2 are still present when Bud8 arrives, and we assume that Bud8 is delivered in vesicles (Figure 8). Finally, all components except Bud8 and Bud5 depart and at some point Cdc10 returns. The appearance of Bud8 at spore polarity sites but the frequent failure to use those sites for polarization in cells lacking *BUD2*, *BUD5*, or *RSR1* fits well with a model in which Bud8 serves an anchor/landmark function in the membrane but itself cannot drive polarization. Considering that our assay for assembly of the site involves Bud8 localization and our assay for functionality of the site involves its usage upon germination, from our data we cannot determine whether in the mutant cells the defect occurs during assembly of the site and/or when cells attempt to use it.

In addition to the Ras-family GTPase Rsr1, other small GTPases function in yeast polarity. Another essential Rho-family GTPase besides Cdc42 is Rho1, which also regulates actin dynamics, particularly under stress conditions (Dong *et al*., 2003), and is required for clathrin-independent endocytosis in yeast (Prosser *et al*., 2011). Rho1 was previously found at the site of spore polarization 90 minutes after germination; no data was provided for mature spores prior to germination (Kono *et al*., 2005). Our studies focused on canonical polarity establishment involving Cdc42, but our data do not exclude a role for Rho1 and its effectors.

It is not at all clear what happens at the polarity site to trigger the (re-)recruitment of Cdc10. Intriguingly, near the end of sporulation Cdc10 undergoes serine phosphorylation at two sites (Wettstein *et al*., 2024). One of these residues, Ser312, is known to be phosphorylated by Cla4 (Versele and Thorner, 2004), an effector downstream of Cdc42 involved in septin assembly in vegetative cells (Kadota *et al*., 2004; Versele and Thorner, 2004). The equivalent site in the Cdc10 homolog in the fission yeast *Schizosaccharomyces pombe* is also phosphorylated specifically during late sporulation (Sivakova *et al*., 2024). In *S. cerevisiae*, Cla4-mediated phosphorylation of Ser312 in late sporulation may increase the affinity of Cdc10 for some factor that remains at the spore polarity site, driving Cdc10 return there. While we did not observe any discrete localization of GFP-tagged Cla4 in sporulating cells (Supplemental Figure 3C), spores lacking *CLA4* show abnormalities in actin organization and morphogenesis upon germination (Kono *et al*., 2005).

### What marks the polarity site after meiotic outer plaque detachment from the membrane?

In addition to the unknown factor(s) that re-recruit Cdc10 to the spore polarity site, we do not know what targets exocyst and Myo2 to a specific membrane site after PSM closure. For the ∼45 minutes that follows PSM closure we have no marker for the polarity assembly site. Our current data do not exclude the possibility that the polarity sites instead form at the locations of PSM closure. However, since the polarity sites are at the ascus periphery, in such a model spores would have to undergo orchestrated rotation within the ascus, for which we see no evidence. These new data are thus consistent with our original model that polarity factors persist at the PSM origin.

The ability of Bud8 to anchor the site to a single, stable location on the cell cortex likely requires interaction of Bud8’s extracellular domain with the inner layer(s) of the spore wall. Indeed, the timing of Bud8 appearance may be programmed to coincide with spore wall biogenesis. What prevents diffusion of the polarity assembly site in the membrane before Bud8 arrives? Clustering of GTP-bound Cdc42 in vegetative cells limits its diffusion at the cortex (Sartorel *et al*., 2018), but this cannot explain the time period before Cdc42 arrives. In vegetative cells the phospholipid phosphatidylserine (PS) is delivered in secretory vesicles and concentrates at sites of polarized exocytosis, where it binds Cdc42 and contributes to the limited diffusion of Cdc42 clusters (Fairn *et al*., 2011; Sartorel *et al*., 2018). Following PSM closure, PS localizes to a single focus per spore (Parodi *et al*., 2015), highly reminiscent of the spore polarity site. However, PS is found throughout the growing PSM, not solely at the PSM origin.

Phosphatidylinositol-4,5-*bis*-phosphate binds exocyst (He *et al*., 2007) and is enriched in the membranes of mature spores (Rudge *et al*., 2004), but in the period following PSM closure a widely-used fluorescent reporter (derived from the pleckstrin homology domain of human phospholipase C δ1 (Stefan *et al*., 2002)) shows signal only around the plasma membrane of the diploid mother cell (Supplemental Figure 3D). A caveat of such protein-based lipid reporters is that they could in principle be outcompeted for binding to their target by a native protein with higher affinity for the same lipid.

### Dispensability of filamentous actin in the late phase of polarity site assembly

Our experiments with *cdc42(G142S)* mutants and latrunculin suggest that filamentous actin is important but not essential for spore polarity site assembly. A requirement for actin filaments in the first phases of assembly fits with the idea that Myo2 transports exocyst along actin cables, which are difficult to detect in living cells (Wirshing and Goode, 2024), though it could also reflect a requirement for endocytosis.

Once delivery of Bud8 begins, however, actin filaments appear to be dispensable for persistence of exocyst and the additional accumulation of Bud8 in the later phases of spore polarity site assembly. A compelling precedent for actin-independent polarization in yeast comes from a study of haploid cells exiting from starvation-induced quiescence, in which actin filaments naturally collapse yet polarized morphogenesis occurs due to random diffusion of secretory vesicles that eventually dock and fuse at the distal pole, presumably binding to polarity factors persisting there (Sahin *et al*., 2008). In this way, a small amount of Bud8 and associated polarity machinery delivered via actin cables could act as a stable docking site for undirected vesicles containing more Bud8 (Figure 8). Indeed, in vegetative cells numerous polarity proteins (Ayscough *et al*., 1997) as well as components of late Golgi vesicles (Rossanese *et al*., 2001) are able to polarize in the presence of Lat A (*i.e.*, in the absence of actin filaments). In fission yeast polarity, actin-independent exocyst localization also operates in parallel with long-range cytoskeletal transport, both downstream of Cdc42 (Bendezú and Martin, 2011).

### Tag-induced exocyst defects and Bud8 trafficking

To monitor exocyst localization, we used GFP fusions to the C termini of exocyst subunits and were fortunate to find subtle defects that revealed functional links between exocyst and the Bud8-marked spore polarity site. C-terminal tags on mammalian exocyst subunits have also been shown to cause functional defects ranging from subtle (Pereira *et al*., 2023) to severe (Ahmed *et al*., 2018), but to our knowledge, this is the first report of a defect caused by GFP-tagging yeast exocyst components. All the various tagged exocyst subunits displayed similar localization dynamics (Figure 2C,D and Supplemental Figure 1D) and in no case did we detect overt defects in sporulation *per se*. We interpret these observations with tagged exocyst as evidence of temporally distinct stages of vesicle trafficking operating sequentially within the process of spore polarity establishment. Others have proposed that yeast cells evolved specific ways to control the timing of exocyst-mediated vesicle fusion during PSM biogenesis (Knop *et al*., 2005). In vegetative cells, an analogous staging mechanism controls the timing of Bud8 translation and/or vesicle delivery: *BUD8* transcription peaks in G2/M, when general vesicle trafficking is targeted to the bud neck, but Bud8 is only delivered to the membrane in the following S phase (Schenkman *et al*., 2002), when trafficking targets the bud tip. Strikingly, we found that tagging the exocyst protein Sec8 caused mislocalization of Bud8 exclusively to bud necks (Supplemental Figure 3E). Mutants lacking the vacuolar SNARE proteins Vam3 or Vam8 exhibit comparable vegetative Bud8 mislocalization (Ni and Snyder, 2001). Tagging exocyst may similarly misdirect otherwise vacuole-bound Bud8 vesicles during sporulation. We think it is unlikely that the appearance of Bud8 signal on all four PSMs in an ascus with tagged exocyst results from normal timing of Bud8 trafficking but delayed PSM biogenesis/closure, because we saw no evidence of such a delay (Supplemental Movie S6).

### Comparisons with other organisms

Although only diploid vegetative cells use Bud8 as a landmark for bud site selection, vegetative haploid cells also deposit Bud8 at their distal poles. Whereas diploid cells can undergo meiosis and sporulation in nutrient-limiting conditions, haploid cells of some strain backgrounds are able to enact a program of highly polarized proliferation that involves repeated budding at the distal pole in a Bud8-dependent manner (Cullen and Sprague, 2002). This pattern of proliferation leads to the invasive dispersal of the population by promoting a linear, rather than radial expansion pattern, superficially reminiscent of the growth habit of filamentous fungi. However, Bud8 homologs are absent from various filamentous fungi (e.g. *Aspergillus* spp., germinating spores of which undergo polarized morphogenesis when they form germ tubes (Harris, 1997)) as well as at least some species that have both a yeast and a hyphal form (*e.g. Candida albicans*) (Seiler and Justa-Schuch, 2010). In *Cryptococcus neoformans*, upregulation of Bud8 is associated with increased virulence (Movahed *et al*., 2015), but mechanistic details are unknown. In the yeast *Yarrowia lipolytica*, the Bud8 homolog localizes all around the cortex of the growing bud but is important for cell separation, not bud site selection (Qing *et al*., 2019). Given the lack of any obvious shared function, the common role for Bud8 during evolution may be as a stable cortical mark that can be repurposed for various cellular tasks.

Beyond the fungi, others have noted similarities between the PSM and the acrosome, a double-bilayer membrane structure in developing spermatocytes that emerges from post-Golgi vesicles and grows to engulf each haploid nucleus (Caudron and Barral, 2009). The similarities extend further: a mature acrosome develops via fusion of proacrosomal vesicles at a single site on the nuclear surface (Khawar *et al*., 2019). Flagellum formation involves plasma membrane morphogenesis from a specific region of the cortex of the developing spermatocyte, conceptually analogous to the polarized morphogenesis from a single site that precedes fusion of yeast gametes. In both cases, septin proteins mark the cortical location from which polarized morphogenesis occurs (Lin *et al*., 2011; Kuo *et al*., 2013).

Moreover, Cdc42 co-localizes with septins during mammalian spermiogenesis (Huang *et al*., 2018).

Spermatocytes are not the only mammalian gametes with similarities to the mechanisms underlying yeast spore polarity. Mammalian oocytes remain dormant for months or years. During oogenesis centriolin, a protein associated with the spindle poles and the mammalian homolog of a yeast SPB component (Nud1), stops contributing to microtubule organization and instead is required for the asymmetry of the divisions that generate the haploid gamete (Sun *et al*., 2017). The Nud1 homology domain of centriolin directly interacts with exocyst (Gromley *et al*., 2005). Further studies of the interplay between polarity proteins and the biogenesis and subsequent polarized morphogenesis of yeast gametes may provide valuable insights into gametogenesis in other organisms.

## MATERIALS AND METHODS

### Yeast strains

*S. cerevisiae* strains (Table S1) were cultured using standard techniques (Amberg *et al*., 2005).

Cells were cultured in liquid or solid (2% agar) rich (YPD: 1% yeast extract, 2% peptone, 2% glucose), synthetic medium (SC; per liter, 20 g glucose or lactose, 1.7 g yeast nitrogen base without amino acids or ammonium sulfate, 5 g ammonium sulfate, 0.05 g tyrosine, 0.01 g arginine, 0.05 g aspartate, 0.05 g phenylalanine, 0.05 g proline, 0.05 g serine, 0.1 g threonine, 0.05 g valine, 0.05 g histidine, 0.1 g uracil, and 0.1 g leucine), or sporulation medium (1% potassium acetate, 0.05% glucose, 20 mg/L leucine, 40 mg/L uracil). Where appropriate to maintain plasmid selection, synthetic medium lacked specific components (uracil and/or leucine).

### Plasmids

*E. coli* DH5⍺ was used for plasmid propagation. Plasmids were introduced into yeast using the Frozen-EZ Yeast Transformation II Kit (#T2001, Zymo Research). To mark PSMs we used pRS426-R20 (Nakanishi *et al*., 2007), which is marked with *URA3* and encodes an RFP-tagged fragment of Spo20 from the *TEF2* promoter and was a gift from Aaron Neiman. We also created a *LEU2*-marked version of this plasmid, called pRS425-R20, by co-transforming pRS426-R20 into yeast cells along with a *LEU2* PCR product made with primers 3_pRS (ACAATTTCCTGATGCGGTATTTTCTCCTTACGCATCTGTGCGGTATTTCACACCG) and 5_pRS (GTGTCGGGGCTGGCTTAACTATGCGGCATCAGAGCAGATTGTACTGAGAGTGCAC) and template plasmid pRS305 (Sikorski and Hieter, 1989). Some yeast strains carried an integrated version of the same Spo20 fragment fused to mKate2 (see Table S1). pFA6a-His3MX6 (Longtine *et al*., 1998) was used as a template for PCR to delete the *BUD5* ORF in some yeast strains (see Table S1). pFA6a- mNeonGreen-SpHis5 was a gift from Jeff Moore (Estrem and Moore, 2019). DLB2968 (*BEM1-GFP*) (Kozubowski *et al*., 2008) and DLB3609 (*GFP-CDC42*) (Wu *et al*., 2015) were gifts from Danny Lew. pFA6a-link-tdTomato-SpHis5 was a gift from Kurt Thorn (Addgene plasmid # 44640; http://n2t.net/addgene:44640 ; RRID:Addgene_44640). pFA6a-link-GFPEnvy-SpHis5 was a gift from Linda Huang (Addgene plasmid # 60782; http://n2t.net/addgene:60782 ; RRID:Addgene_60782). pHP1109 (*RAX1-GFP*) (Kang *et al*., 2004a) and pHP2274 (*mNeonGreen-BUD8*) (Kang *et al*., 2022) were gifts from Hay-Oak Park. Lifeact-GFP was expressed from plasmid p415-Cyc1-Lifeact-GFP (Lin *et al*., 2010), a gift from Jeff Moore. Lifeact-3xmNG was expressed from plasmid pBG2654 (Wirshing and Goode, 2024). pRS426GFP-2xPH(PLCδ) was used as a reporter for phosphatidylinositol-4,5-*bis*- phosphate (Stefan *et al*., 2002). We created integrating plasmids for *mScarlet3-BUD8* (named G01288) and *mScarlet2-BUD8* (named G01287) via *in vivo* recombineering in *E. coli* (Kostylev *et al*., 2015).

Briefly, we designed, purchased (Twist Bioscience) and PCR amplified synthetic sequences encoding mScarlet3 or mScarlet2 flanked by sequences homologous to the sites where digest with *Not*I excised the mNeonGreen sequence from pHP2274. These PCR products were co-transformed with the digested pHP2274 into DH5⍺. Full sequences of these plasmids were determined and are available upon request. For CRISPR-Cas9-mediated introduction of the *cdc42(G142S)* mutation, we used the MyLo method (Bean *et al*., 2022) with annealed oligos encoding a gRNA (CGUCGGUACGCAGAUUGAUCUA) with *Bsa*I-digested plasmid pBBK05 (a gift from Vincent Martin, Addgene plasmid # 178989; http://n2t.net/addgene:178989; RRID:Addgene_178989).

### Culture growth, sporulation, and germination

Liquid YPD cultures were inoculated with cells from YPD or synthetic selective plates and grown at 30°C overnight. Aliquots of cultures were pelleted at 1,000 x *g* for 2 min and resuspended in 5x the volume of sporulation medium. Imaging of sporulating cells was performed after 24 h in sporulation medium. To induce germination, cells incubated in sporulation medium for at least 7 days were diluted into synthetic medium for imaging.

### Microscopy

Concanavalin A was used to coat the bottom of Cellview Cell Culture Slides (#543078, Greiner Bio-One; Kremsmünster, Austria). Excess concanavalin A was removed and wells were washed with sporulation or synthetic medium. Fresh medium was then added to each well before the addition of cells. Immediately after, the slides were centrifuged at 250 x *g* for 2 min at room temperature and then used for imaging. Sporulating cells and mature spores were imaged at room temperature while germinating spores were imaged at 30°C. Z-series covering a 10 µm range were acquired using a 0.5 µm step size at 15-min intervals for varying lengths of time depending on the experiment. For Figure 1B and D, Figure 6C, and Video S6, images were collected on a spinning disk confocal Nikon Ti-E microscope equipped with a 1.45 NA 100× CFI Plan Apo objective, piezo electric stage (Physik Instrumente; Auburn, MA), spinning disk confocal scanner unit (CSU10; Yokogawa), 488-nm laser (Agilent Technologies; Santa Clara, CA), TI-LA FRAP module, and an EMCCD camera (iXon Ultra 897; Andor Technology; Belfast, UK) using NIS Elements software (Nikon).

All other images were collected on a Nikon Ti2-E inverted microscope equipped with 1.45 NA 100x CFI Plan Apo objective (Nikon Inc; Melville, NY), Nikon motorized stage, Prior NanoScan SP 600 mm nano- positioning piezo sample scanner (Prior Scientific; Rockland, MA), CSU-W1 T1 Super-Resolution spinning disk confocal, SoRa disk with 1x/2.8x/4x mag changers (3i; Denver, CO), 488 nm, 560 nm, and 647 nm laser, and a prime 95B back illuminated sCMOS camera (Teledyne Photometrics; Tuscon, AZ).

Fiji was used to process and analyze images (Schindelin *et al*., 2012). Image stacks were first processed as maximum projections before downstream analysis. To stabilize time course images for analysis, Fast4dreg (Laine *et al*., 2019) was used stabilize time course images for analysis. The “time estimate+apply” function was used to correct for X and Y drift using the transmitted light or red channel. These settings were then used to correct drift in the other channels with the “time apply” function.

Corrected channels were then merged for cropping. Particle tracking was performed using Trackmate (Ershov *et al*., 2022). Puncta identification was performed on z-projected images of three evenly spaced images centered around puncta present at t=0, spanning a 1.5 µm distance. A quality score of >0.02 was used as an initial filter, followed by additional filtering to only include tracks with <1 gap that appeared in >11 of 13 total timepoints. The following settings were used: LoG detector with an estimated object diameter of 2 µm and a quality threshold of 0 using sub-pixel localization. Simple LAP tracker was used with a linking max distance of 1 µm, gap-closing max distance of 1 µm, and gap-closing max frame gap of 2. From these tracks, we selected only those that, by eye, appeared to be unambiguous puncta in tetrads. For Supplemental Video S1, bleach correction was applied using histogram matching (Miura, 2020).

### Latrunculin Treatment

Latrunculin A (#AGCN20027C500, AdipoGen Life Sciences) and B (#10010631, Cayman Chemical) were dissolved in DMSO to create a stock concentration of 20 mM. 1 µL of latrunculin or DMSO was added to a 100-µL cell suspension (200 µM working concentration) before immediate transfer to wells for adherence and imaging.

## ACKNOWLEDGEMENTS

We thank Michael Knop (University of Heidelberg) and Danny Lew (Massachusetts Institute of Technology) for valuable feedback on the manuscript, and Jeff Moore (University of Colorado Anschutz Medical Campus School of Medicine) for generous access to microscopes. This work was supported by funds from the National Science Foundation, grant number 1928900, and from the National Institute of General Medical Sciences, grant number R35GM148198.

## Abbreviations

GAP: GTPase activating protein
GEF: guanine nucleotide exchange factor
MOP: meiotic outer plaque
PS: phosphatidylserine
PSM: prospore membrane
SPB: spindle pole body

