## Supplemental Figures 1-3 for "Roles for the canonical polarity machinery in the *de novo* establishment of polarity in budding yeast spores"

SUPPLEMENTAL MATERIALS

Cooperman and McMurray

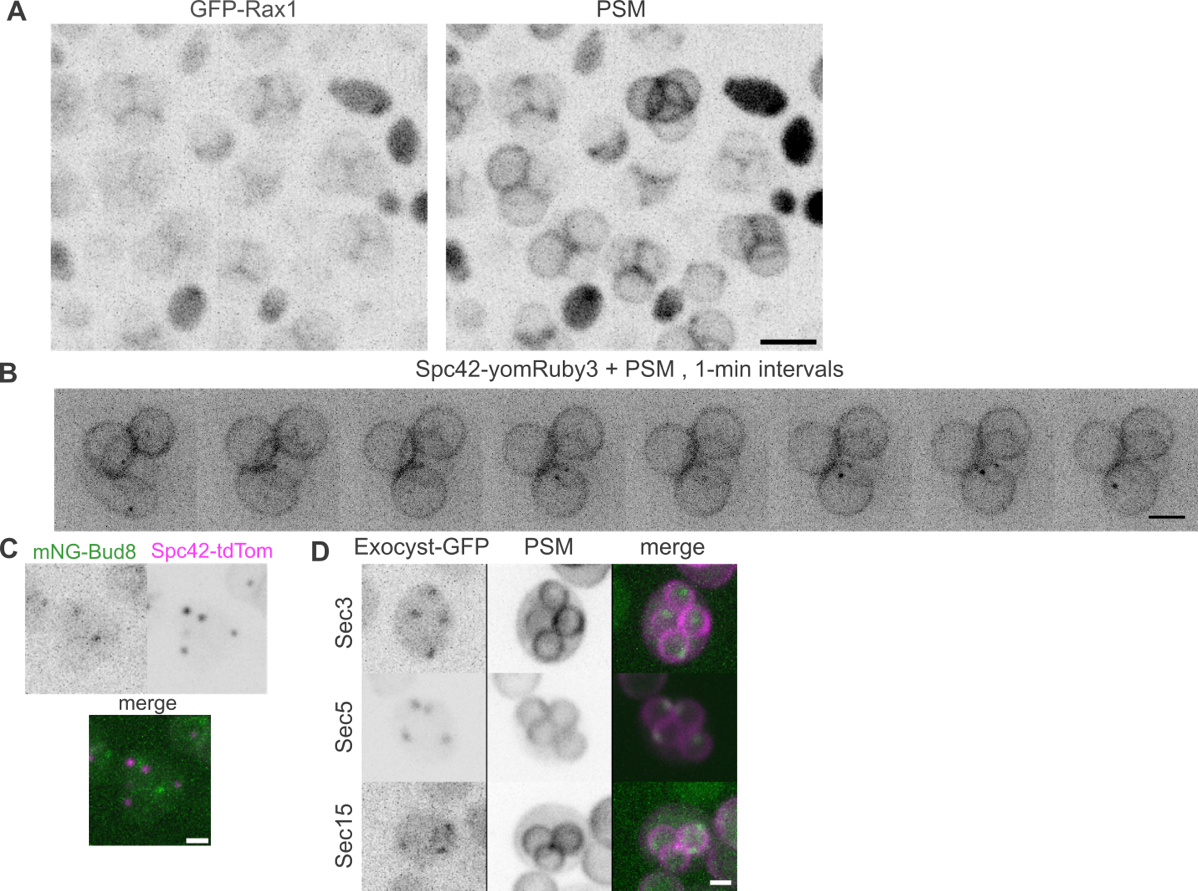

**Supplemental Figure 1. Landmark protein, exocyst, and spindle pole body localization during sporulation**

(A) Mature asci (and some diploid cells that failed to sporulate) of strain 3F245D63 carrying plasmid pRS426-R20, expressing GFP-Rax1 and the RFP-tagged PSM reporter. Scale bar, 5 µm.

(B) A representative ascus of strain FYBY7483 co-expressing two proteins tagged with red fluorescent proteins, the spindle pole body component Spc42 and the PSM reporter (pRS425-R20). Images were captured at 1-min intervals starting just before PSM closure. Scale bar, 2 µm.

(C) A representative mature ascus of strain H07266 co-expressing tdTomato-tagged Spc42 (“Spc42-tdTom”) and mNeonGreen-tagged Bud8 (“mNG-Bud8”). Scale bar, 2 µm.

(D) Cells expressing the RFP-tagged PSM reporter (from pRS425-R20) and GFP-tagged versions of the indicated exocyst subunits were imaged while undergoing sporulation. Strains were H07191, H06741, and H07182. Scale bar, 2 µm.

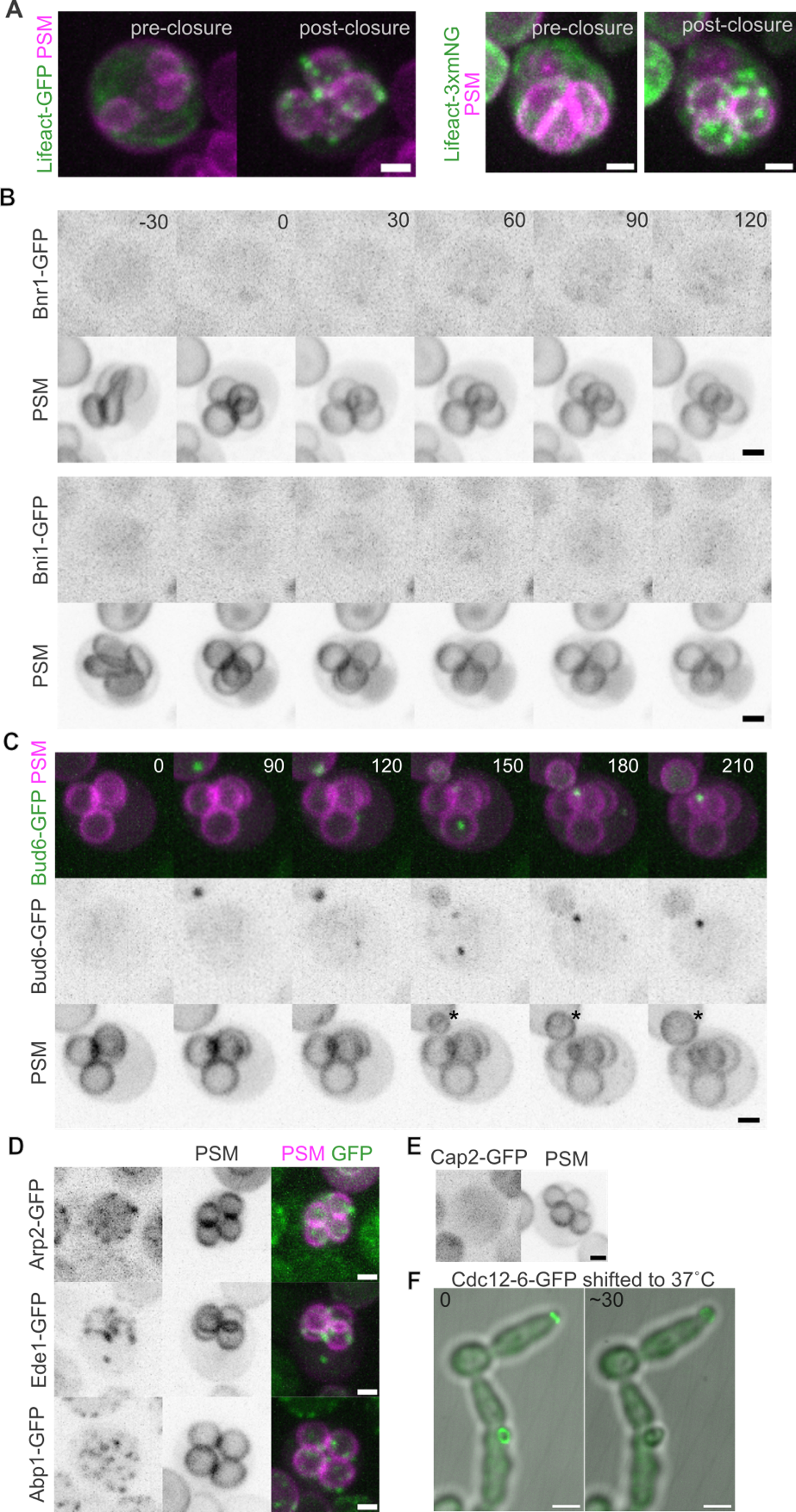

**Supplemental Figure 2. Testing requirements for the actin cytoskeleton in assembly of the spore polarity site.**

(A) Filamentous actin was visualized in cells undergoing sporulation using Lifeact-GFP. Representative images show cells of strain D2191750 carrying plasmid p415-Cyc1-Lifeact-GFP (left) or of strain FYBY7483 carrying plasmid pBG265 (right) before and after PSM closure. Scale bar, 2 µm.

(B) The actin-nucleating formin proteins Bnr1 (top) or Bni1 (bottom) were tagged with GFP and imaged in cells carrying plasmid pRS425-R20 undergoing sporulation. Representative asci of strains D07B95E7 or 48C7D450 are shown. Scale bars, 2 µm.

(C) A representative ascus of strain 84ED8927 carrying plasmid pRS425-R20 expressing Bud6-GFP and the RFP-tagged PSM reporter undergoing sporulation. Numbers indicate minutes elapsed since PSM closure. Scale bar, 2 µm. The asterisk indicates a bud produced by an adjacent vegetative cell.

(D) Cells co-expressing the PSM reporter and GFP-tagged versions of the indicated markers of endocytosis were imaged during sporulation. Shown are representative asci of strains 786C0E58, 6C7CA06E, and BD1686E4, carrying plasmid pRS425-R20. Scale bars, 2 µm.

(E) A Cap2-GFP-expressing ascus of strain A76EDCB8 carrying plasmid pRS425-R20 was exposed to 200 µM Latrunculin B. Scale bar, 2 µm.

(F) Cells of strain GFY-1086 were cultured at 30˚C in rich medium then transferred to complete synthetic liquid medium in a multi-well imaging chamber. A cell was imaged before and ~30 minutes after the temperature of the microscope stage was raised to 37˚C.

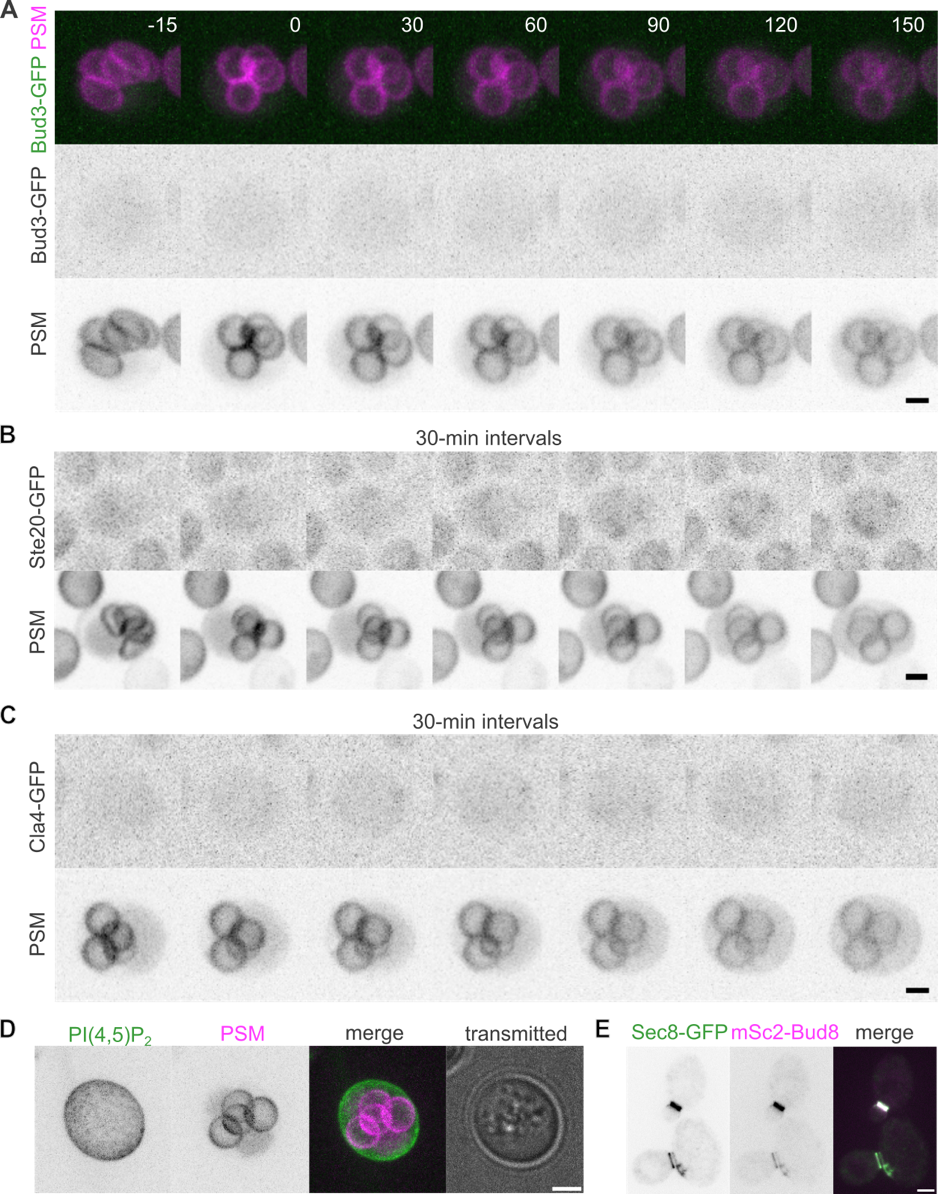

**Supplemental Figure 3. Localization patterns of additional polarity-related factors.**

(A) A representative ascus of strain H07162 carrying plasmid pRS425-R20 expressing Bud3-GFP and the RFP-tagged PSM reporter undergoing sporulation. Numbers indicate minutes elapsed since PSM closure. Scale bar, 2 µm.

(B) A representative ascus of strain 1644910E carrying plasmid pRS425-R20 expressing Ste20-GFP and the RFP-tagged PSM reporter undergoing sporulation. Scale bar, 2 µm.

(C) A representative ascus of strain 5594A743 carrying plasmid pRS425-R20 expressing Cla4-GFP and the RFP-tagged PSM reporter undergoing sporulation. Scale bar, 2 µm.

(D) A representative ascus of strain FYBY7483 expressing a GFP-tagged reporter of PI(4,5)P_2_ (pRS426GFP-2×PH(PLCδ)) and the RFP-tagged PSM reporter (from plasmid pRS425-R20) imaged shortly following PSM closure. Scale bar, 2 µm.

(E) Two representative budding cells of strain H07266 co-expressing GFP-tagged Sec8 and Bud8 tagged with mScarlet2 (“mSc2”). Scale bar, 2 µm.
